# Pooled functional genomic screens for intracellular calcium effectors

**DOI:** 10.1101/434910

**Authors:** Anjali Rao, Joel M. Kralj

## Abstract

Cytoplasmic calcium transients relay cellular signals on timescales from milliseconds to hours, and the dynamic nature of calcium signals has slowed functional genomic screening for cellular calcium effectors. Here, we present a new strategy to identify calcium handling genes via a pooled knockdown employing the calcium-sensitive photo-switchable fluorescent protein, CaMPARI. This assay, cal-Seq, enabled identification of regulators for both cellular desensitization and histamine induced calcium signaling including GPR99, a leukotriene binding receptor, with implications in asthma treatment.

---

Calcium is a critical second messenger, and intracellular calcium levels are precisely controlled by numerous cellular effectors^1^. Basal cytoplasmic calcium levels are low, but can be increased within milliseconds to modulate cellular behavior. Calcium dynamics encode cellular signals through modulation of amplitude, frequency, and spatial location of the concentration flux, and numerous diseases are linked to mutations in calcium handling proteins^2^. Despite this critical importance, the transient nature of cytoplasmic calcium has slowed discovery of calcium effectors due to the difficulties of screening transients at genomic scales.

Genome-wide screens (GWS) employing pooled shRNA and CRISPR technologies are powerful tools for gene function discovery^3^. However, current technology limits pooled screens to probing slow physiological processes like cell survival, proliferation, or gene expression. Rapid cellular changes, like calcium transients, are too short-lived to screen and sort entire libraries, and require imaging-based screens of individual genes in an arrayed format^4,5^. However, arrayed screens require long experiment times and the use of expensive robotic liquid handling and imaging. Conversely, pooled genomic screens can be economical and fast.

In this paper, we developed a new strategy to profile calcium handling genes in a pooled format using CaMPARI, a unique, genetically encoded, fluorescent calcium sensor^6^. CaMPARI photoconverts from green to red fluorescence only in the presence of both Ca^++^ and 405 nm light. The irreversible photoconversion acts as a mechanism to “lock-in” the magnitude of the calcium signal (red to green fluorescence ratio) at a specific, user defined instant in time. The calcium concentration can be measured post hoc with a microscope or Fluorescence Activated Cell Sorting (FACS) machine. We coupled CaMPARI with the scalability of pooled genome-wide knockdown libraries to develop a new functional, high-through assay to rapidly map genes that regulate intracellular calcium levels (Fig. 1A). The protocol started with a pool of isogenic, CaMPARI expressing cells treated with lentivirus containing a pooled knockdown library. Chemical stimulation triggered calcium influx, followed by exposure to 405 nm light to photoconvert cells with high cytoplasmic calcium from green to red fluorescence. All cells are exposed to identical chemical conditions, and the number of measured cells is > 100x library coverage, both of which help minimize the effects of population heterogeneity. Green cells (low calcium) are sorted via FACS, and the associated genetic perturbation (shRNA) sequences are read via the Illumina platform. Genes enriched among green cells are then identified by computational analyses, and top hits and pathways are validated individually. The entire screen, which we termed cal-Seq, can be performed in triplicate in 4 weeks using instruments available to most researchers.

**Figure 1:**
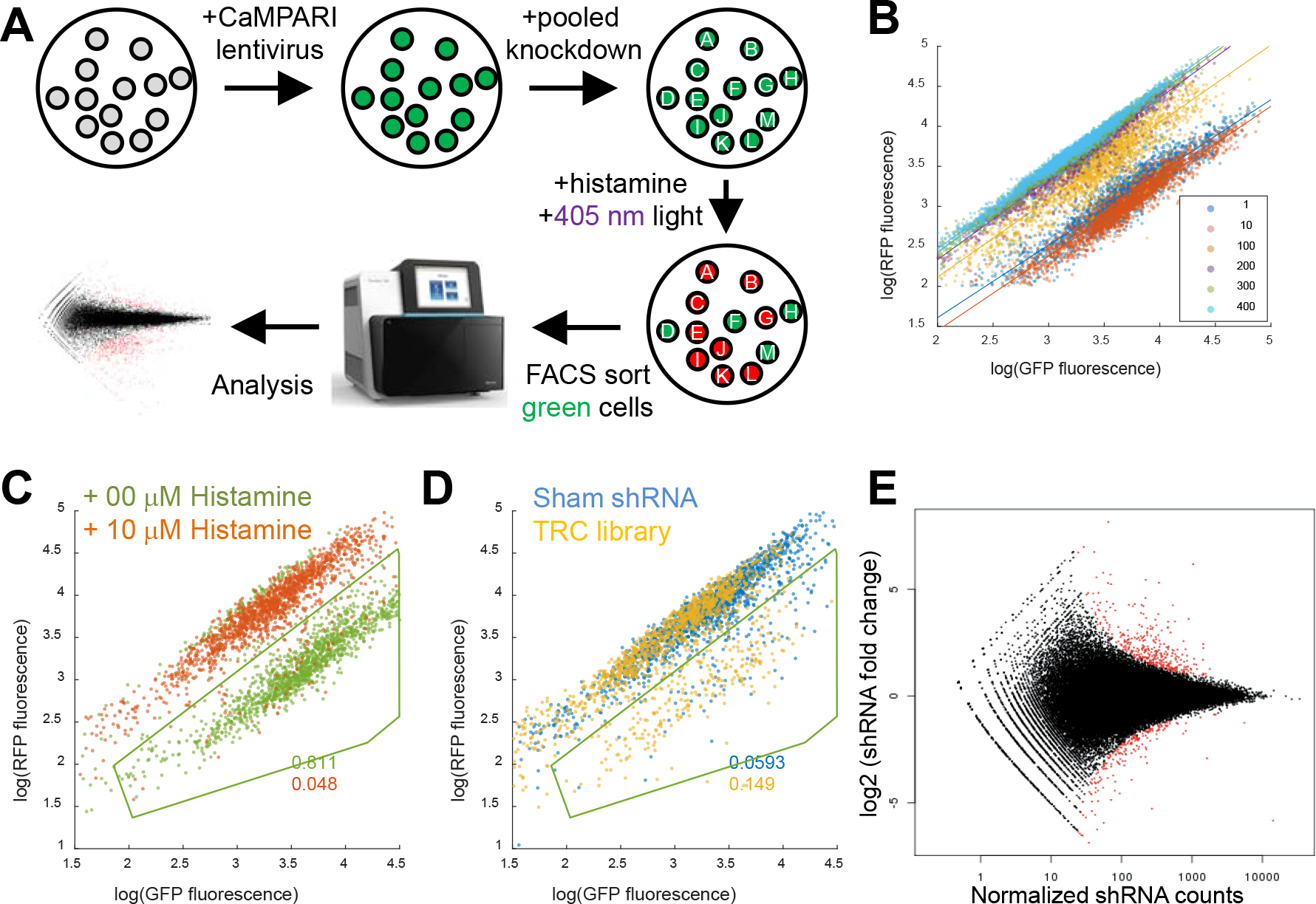
Implementation and proof of concept for cal-Seq assay. (A) Schematic for a pooled functional genomic assay combining CaMPARI, FACS, and next-gen sequencing. (B) Measured Kd for CaMPARI photoswitch using the ratio of red to green fluorescence. Varying amounts of cytoplasmic calcium were photoconverted under otherwise identical conditions leading to increased red fluorescence. The Kd was measured to be 124 nM. (C) Histamine stimulation (10 μM, red dots) caused a decrease in the fraction of green cells as compared to addition of basal medium (0 μM, green dots) from a manually selected ROI. (D) HeLa cells infected with the TRC1 shRNA library (yellow dots) showed an increased fraction of green cells as compared to a sham shRNA under identical chemical and illumination conditions. (E) MA plot of genes enriched in the green population of cells performed in triplicate. Red dots indicate genes with a p-adjusted value < 0.05.

As proof of concept, we used this assay to identify genes essential for histamine-induced calcium in HeLa cells. CaMPARI acted as both a photoswitchable and real-time calcium indicator upon histamine stimulation, as previously reported^6^ (Fig 1B, S1). Fluorescence cytometry experiments measured a photoswitch Kd of 124 nM Ca^++^ (Fig 1B, S2), similar to imaging measurements^6^. The histamine induced calcium combined with 405 nm light (40 seconds) showed a green to red fluorescence conversion as indicated by cytometry (Fig 1C and S2). Addition of a pooled lentiviral library (TRC1/1.5 shRNA^7^) increased the fraction of green cells (low calcium) ~9% as compared to scrambled shRNA infected cells (Fig.1D). This population of green cells was sorted via FACS, the associated shRNAs were identified using the NextSeq platform (Fig S3), and enrichment was quantified from 3 biological replicates with DeSeq^8^.

DeSeq analysis identified 348 shRNAs enriched in the green population (p-adjusted < .05, Fig 1E, Table S1). We selected 23 hits with GO terms related to calcium handling to individually knock down in CaMPARI-expressing HeLa cells. Measured via cytometry, 29% significantly increased the fraction of green cells (Fig.S4). We attributed this low significance to low signal-to-noise ratio (SNR) CaMPARI cytometry measurements, as well as common issues with functional genomic screens^9^. These gene knockdowns were also measured with Twitch2B^10^, a real time, ratiometric calcium indicator. Using this assay, 65% of knockdowns showed significantly reduced calcium influx compared to a sham knockdown. Hence, cal-Seq was successful in identifying genes important for calcium handling.

Among the top hits were several expected genes and pathways. Pathway analysis revealed known regulators of InsP3R, an essential downstream component of histamine signaling, including isoforms of PKC (PRKCA, PRKCQ), PKA (PRKAR1a, PRKX), CamK2A, and Fyn^11^ (Fig.S4, S5), as well as regulators of inositol-3-phosphate (ITPKC, PIP5K1B, SKIP, PIK3AP1, PIK3R1). GBγ and Gαi signaling pathways were enriched, as well as mediators of NFAT/calcineurin signaling^12^. PPIF, a calcineurin inhibitor, and proteins known to be involved in axonal guidance like EphA10, Slit1, and Sema4F also appeared. These proteins are known to affect calcium signaling^13^, and are involved in inflammation^14^, a key function of histamine. Surprisingly, H1R was not a hit. Only a single shRNA (TRC-62) modestly reduced H1R expression and CaMPARI photoconversion (Fig.S6) in our hands. However, chemical inhibition of H1R and PKC significantly increased the percent of green cells (Supp Fig 6]). These data highlight the need for new sensors optimized for FACS screening to improve the SNR of the cal-Seq data.

GPR99 (OXGR1), a GPCR receptor that is proposed to act as a leukotriene receptor, appeared as one of the top hits, significantly decreased calcium in both cytometry and real-time measurements, and was investigated further due to a reported role in allergic inflammation^15,16^. HeLa cells infected with GPR99 shRNA showed a 3-fold decrease in GPR99 protein levels as compared to the wild type (Fig.S7). Cytometry and real time measurements showed decreased histamine induced calcium transients as compared to a sham (Fig 2A,B). Over-expression of GPR99 using a plasmid ORF in the knock-down cells did not rescue this phenotype (Fig.S7). The ORF over-expressed protein ran at lower molecular weight than the mature protein at 55 kDa which is thought to be glycosylated^17^. It is likely the exogenously expressed protein is trapped in the endoplasmic reticulum instead of getting glycosylated and transported to the plasma membrane.

**Figure 2:**
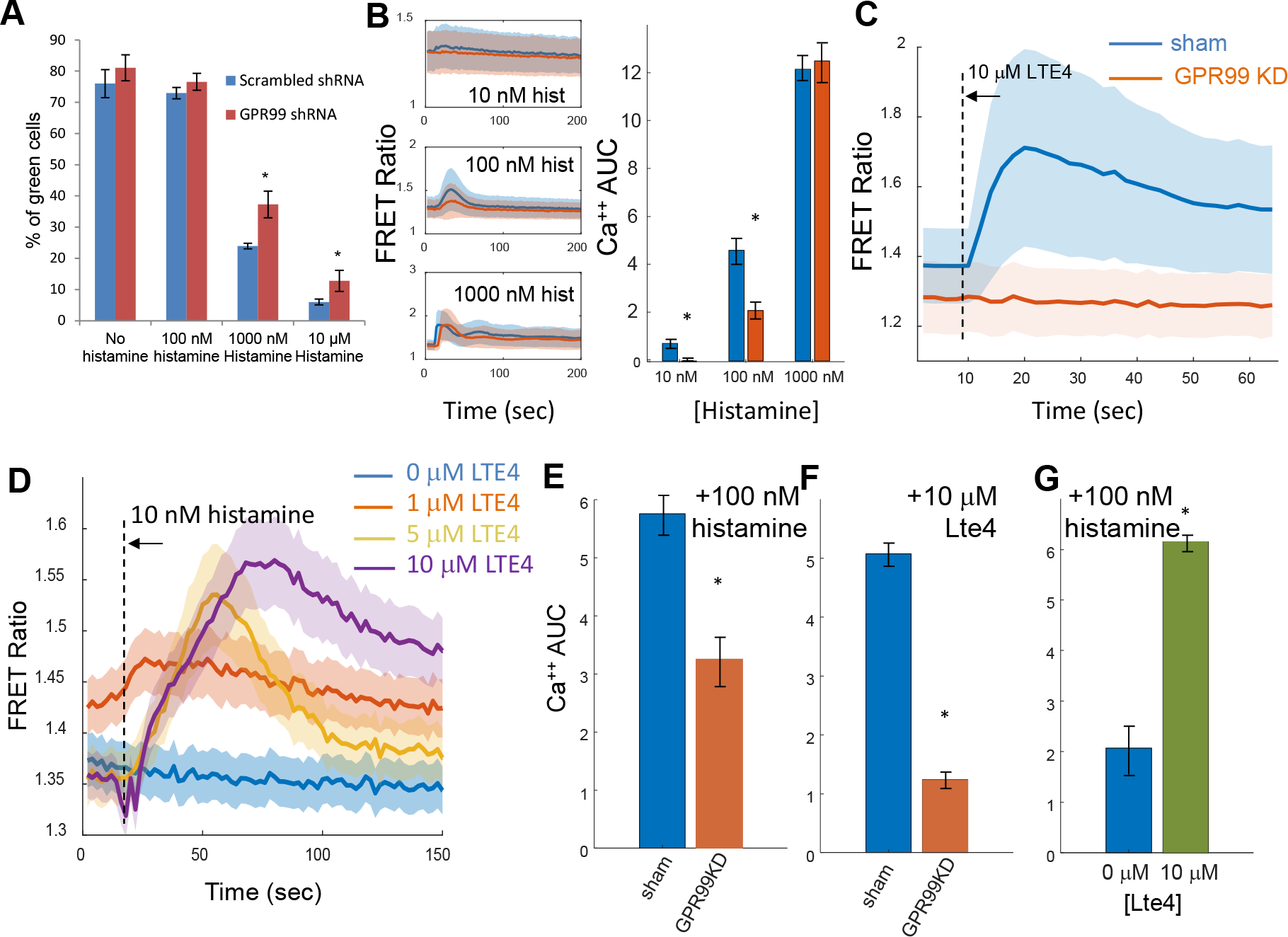
GPR99 regulates histamine induced calcium influx in epithelial cells. (A) Cytometry measurements comparing the fraction of green cells from a sham shRNA (blue) compared to GPR99 knockdown (red) at increasing levels of histamine stimulation. (B) Real-time cytoplasmic calcium measured with Twitch2B. The mean (solid) and standard deviation (shaded) traces of HeLa cells upon stimulation with increasing amounts of histamine. A GPR99 knockdown shRNA (red) had decreased calcium influx as compared to a sham shRNA (blue). Area under the curve (AUC) measurements are shown for the same data by integrating the first 40 seconds after histamine stimulation for the same data.

Leukotriene E4 (Lte4) has been proposed as the ligand for GPR99 based on *in vitro* and knock-out mouse studies^15,16^. Addition of Lte4 to HeLa cells increased cytosolic calcium concentrations dependent on GPR99 expression (Fig. 2C). In WT cells, pretreatment with Lte4 caused an increased calcium response to histamine stimulation that was dependent on the presence of GPR99 (Fig 2D and S8). We further investigated the relationship between GPR99 and histamine signaling in Beas2B cells, a widely used epithelial cell line for modeling upper respiratory tract infections in asthma^19^. Similar to HeLa cells, Beas2B cells showed both histamine and Lte4 induced calcium currents (Figs 2E-G, S9). GPR99 knockdown reduced histamine induced flux and eliminated the Lte4 response. Furthermore, the presence of excess Lte4 increased the histamine induced calcium response. These data show synergy between histamine and Lte4 in epithelial cells and provide molecular context to similar observations in embryonic cancer cells^18^. In the past, synergism between drugs targeting leukotriene receptors and histamine receptors has been proposed based on observations from clinical trials in asthmatic patients^20^ indicating cross-talk between these two pathways. Current drugs on the market target cysLTR1 which have low affinity towards Lte4, a proposed biomarker for allergic inflammation like asthma^21^. GPR99 has higher affinity towards Lte4^15^, and hence, targeting GPR99 might have additional benefits for patients with pathological inflammatory conditions.

To extend the potential utility of cal-Seq, we leveraged the timing control between chemical stimulation and lock-in light pulse to identify effectors of receptor desensitization. Cells exposed to the same stimulus reduce their response from numerous cellular factors including arrestins, endocytosis, and ubiquitination^22^. We applied a stimulation protocol similar to previous reports known to induce homogenous histamine desensitization in HeLa cells^23^ (Fig 3A, S10). Repeated histamine stimulation for 5 minutes followed by washout resulted in an increased fraction of green (desensitized) cells as compared to the initial exposure indicative of desensitization (Fig 3B,C). TRC1 lentivirus application increased the fraction of red cells as compared to a sham knockdown (Fig 3D). The red pool was sorted, sequenced, and analyzed for differential expression in triplicate. DeSeq analysis yielded 140 shRNA clones significantly enriched (p-adjusted < 0.1) in the red fraction, which is lower than the number of hits identified from histamine stimulation (Fig 3E, Table S2). The lower number of significant genes is likely due to a higher background of red cells before library knockdown which increased noise. Despite the decreased SNR, genes previously associated with desensitization were enriched in the red population including arrestins, endocytic regulators, and ubiquitin modifiers and suggested our screen identified relevant proteins (Fig.S11). Real-time imaging confirmed a decreased desensitization in knockdowns of ARRDC4 and TOM1L2 compared to a sham knockdown (Fig.S10). Future work will investigate specific genes found on our list and how they mediate desensitization.

**Figure 3:**
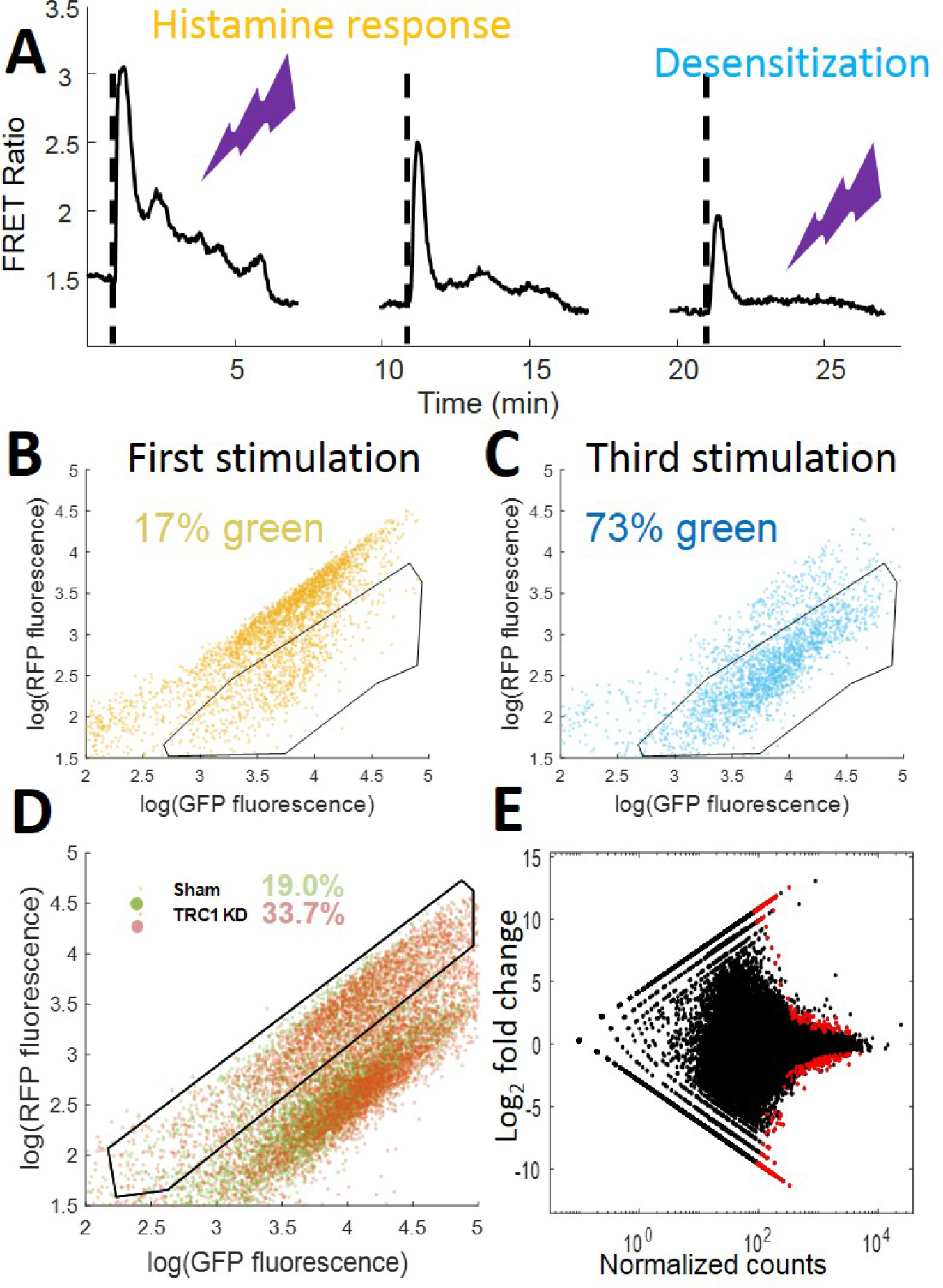
Cal-Seq can be modified to identify genes associated with desensitization. (A) Twitch measurements of HeLa cells upon repeated stimulation and washout of 10 μM histamine (black dashed lines). The purple flashes indicate either 405 nm exposures times for either histamine response (yellow) or desensitization (blue). (B) Cytometry data showing few green cells in a manually selected gate during the first stimulation with histamine. (C) Cytometry data showing increased green cells (desensitized) after the third stimulation. (D) The fraction of red cells increased in a TRC library (red) compared to a sham (green) shRNA infection indicating cells that are deficient in desensitization. (E) MA plot of genes enriched in the green population of cells performed in triplicate. Red dots indicate genes with a p-adjusted value < 0.05.

In conclusion, we believe that cal-Seq can be used to better understand the cellular components giving rise to transient calcium signals in mammalian cells. Our screen identified genes essential for histamine-induced calcium influx and histamine induced desensitization in HeLa cells. Published CaMPARI mutants with variable Kd are available^6^ will enable fine tuning of dynamic range for a variety of experiments in different cell types or organelles. Improved CaMPARI mutants with higher sensitivity, or non-fluorescent calcium integrators, can further improve the sensitivity of this screen. We envision long-term use of this technique to identify potential genes to target with small molecules in disorders with disrupted calcium homeostasis like Alzheimer’s disease, ALS, cardiac conditions and aging.

## Methods

### Plasmids

A gBlock containing CaMPARI (pcDNA3-CaMPARI was a gift from Loren Looger (Addgene plasmid # 60421)) fused with P2A peptide followed by a blasticidin cassette was obtained from Integrated DNA Technologies, Inc. (Coralville, IA). This gBlock was cloned into PmeI and SmaI restriction sites of pWPXL plasmid (pWPXL was a gift from Didier Trono (Addgene plasmid # 12257)) via Gibson cloning. pHuji (pBAD-pHuji was a gift from Robert Campbell (Addgene plasmid # 61555)) was cloned into pLentiCMV plasmid (pLenti-CMV-MCS-GFP-SV-puro was a gift from Paul Odgren (Addgene plasmid # 73582)) using BamHI/SalI via Gibson cloning. Twitch-2B (Twitch-2B pRSETB was a gift from Oliver Griesbeck (Addgene plasmid # 48203)) was cloned into BamHI and EcoRI of pWPXL using restriction digestion and ligation. H2B in a lentiviral plasmid was a gift from the Spencer lab. Human TRC1/1.5 shRNA library (~100,000 shRNAs), individual shRNAs for the hits in the screen, and open reading frame (ORF) plasmid for GPR99 were obtained from Functional Genomic Facility, University of Colorado, Denver.

### Cells

HeLa cells were obtained from ATCC (Manassas, VA), and maintained in DMEM, 10% fetal bovine serum, 2mM L-glutamine, and 500µg/ml penicillin-streptomycin (Thermo Fisher Scientific, Waltham, MA) at 37 °C. Lenti-X 293T cells for lentivirus production were obtained from Clontech (Mountain View, CA), and grown in DMEM10 medium supplemented with 1mM sodium pyruvate and 25mM HEPES (pH 7). Beas2B cells were purchased from ATCC (Manassas, VA), and maintained in LHC-9 media (Thermo Fisher Scientific, Waltham, MA).

### Lentivirus production

60-70% confluent 293T cells in 15 cm cell culture dishes were transfected with 15 µg p∆8.9, 7 µg VsVg, and 11 µg gene of interest using polyethylenimine. Media was changed after 4 hours. Virus particles were harvested from the supernatant 48 hours post-transfection, and filtered through 0.45 µm filter, and stored at −80°C. These viral suspensions were then added directly to cells in 3.5 mm dishes (~1 mL of supernatant), and incubated for 12 hours. Viral preps derived from this method gave us about 90-95% infection efficiency in HeLa cells with no apparent cell death. Antibiotic treatment (3 µg/ml Puromycin (2 days) or 5 µg/ml Blastacidin (7 days)) was started 48 hours post-infection. The cells were allowed to recover in antibiotic-free media for a minimum of 12 hours before experiments were performed.

### Photoconversion and cell sorting

HeLa cells were washed and covered with HBSS/20 mM MOPS. Cells were places in a custom built LED lightbox capable of emitting 405 nm light at 400 mW/cm^2^. The lightbox used 54 405 nm LEDs arrayed inside a reflective chamber (LEDSupply.com, *#*A008-UV400-65).

For direct stimulation of histamine treated cells, histamine (Sigma Aldrich, St. Louis, MO) at appropriate concentration was added onto the cells, and the 405nm light was turned on immediately for 40 seconds. Post-photoconversion the cells were trypsinized and suspended in HBSS/MOPS buffer supplemented with 10 mM EGTA. Subsequently, cells were sorted on the BD FACSAria Fusion (BD Biosciences, San Jose, CA) using 488 and 561 nm lasers. Sorted cells were collected in media, and spun at 400 ×g. Genomic DNA was extracted from the cells using the DNeasy Blood and Tissue Kit (Qiagen, Valencia, CA). Genomic DNA was measured using the NanoDrop.

For the desensitization experiments, histamine was added at 10 μM and left to incubate for 5 minutes. After 5 minutes, all the medium was replaced by histamine free medium for an additional 5 minutes. This process was repeated 3 times. On the third application of histamine, a 405 nm lock in pulse was delivered for 40 seconds, 2.5 minutes after histamine addition. Trypsinization and sorting were identical to the previous experiments.

### Library prep and deep Sequencing

Our library preparation strategy was similar to a previously published study ^24^. Briefly, shRNA cassette was isolated from genomic DNA via PCR. 12 PCR reactions with 600 ng genomic DNA template per reaction were used to get a 10-fold coverage of the unsorted library. The resulting PCR product was digested with XhoI to recover a single strand of shRNA. Barcodes were ligated to the XhoI-digested fragments, and a second round of PCR was performed to include the Illumina adaptor sequences. Resulting products were gel purified, and several quality control tests for the library were done using Bioanalyzer assay, Qubit, and qPCR. Barcodes allowed multiplexing of samples, and pooled samples were loaded at 5 pM (30% PhiX was spiked in) and sequenced on a NextSeq 500 at the Sequencing core facility at Biofrontiers Institute, University of Colorado, Boulder. Sequencing was performed at 500X depth. We obtained 340 million reads, and 95% of these reads were > Q30.

### Sequencing data analysis

The quality of the reads was assessed using the FastQC program run using a Linux interface (Supp Fig. 4). Data preprocessing was performed using the FASTX toolkit. Demultiplexing was performed using FASTX Barcode Splitter, and low quality reads were discarded using the FASTQ Quality Filter. Pre-processed reads were mapped to the shRNA library using the Bowtie aligner. Around 94% of the shRNA clones in the library were detected in our reads suggesting good coverage. Differential reads between our unsorted and sorted samples were estimated using DESeq analysis^8^ in R. Pathway analysis was performed with the Ingenuity Pathway Analysis software package (Thermo-Fischer) by using genes with a p-adjusted value < 0.05.

### RNA analysis

RNA was extracted from cultured cells using Trizol (Life technologies). Total RNA was treated with Turbo DNase (Life technologies) followed by phenol/chloroform extraction. 500 ng of RNA was reverse transcribed using random hexamers.

### Western Blot analysis

Western blots were used to measure the protein concentration of GPR99 in both HeLa and Beas2b cells. Cells were infected with a sham shRNA or a shRNA targeting GPR99, selected with puromycin and incubated for 2 days. Cells were disrupted using RIPA lysis buffer (Thermo-Fischer, #89900) according to the manufacturers directions. 10 μg protein was loaded into each lane of a NuPage 4-12% gel (Thermo-Fischer, #NP0323BOX). Protein was transferred onto a PVDF filter (Thermo-Fischer, #LC2005) and blocking was achieved with 5% BSA for 1 hour at 4 °C. A primary antibody for GPR99 (Abcam, #ab140630) or beta-tubulin was soaked for 2 hours at 4 °C at a dilution of 1:1000. HRP secondary antibodies were used to generate contrast. Quantification was performed in ImageJ (NIH).

### Imaging

Live cell imaging (room temperature) of HeLa cells expressing lentivirus-based Twitch-2B and H2B was carried out using a Nikon Spinning Disc Confocal microscope at the BioFrontiers Advanced Light Microscopy Core, University of Colorado, Boulder. Cells were imaged on tissue culture plastic and drug additions were pipetted during data acquisition. Cells were imaged with a 20X, NA 0.5 objective onto an EMCCD (Ultra888, Andor). Cells were illuminated with 445 nm and 515 nm lasers to excite the CFP and YFP fluorophores on Twitch, respectively. Movies were acquired by taking a 100 ms frame every 2 seconds, with the illumination sources off during the wait. Image segmentation was achieved by using the BFP H2B mark after the histamine stimulation. The cytoplasmic signal was generated by extending a ring of 5 pixels around the nucleus and averaging across those pixels to obtain an intensity for each cell. Intensities were then extracted for each cell, and the entire cell population was used to calculate the mean and standard deviation. The calcium AUC was measured by integrating using the trapezoidal rule from the histamine addition for 40 seconds to mimic the photoconversion.

All image analysis scripts were custom written in Matlab R2017a, and all scripts will be made available to researchers upon request.

## Acknowledgements

We thank Ben Dodd for his inspiration for this project, and Giancarlo Bruni, Andrew Weekley, Humza Ashraf, and Ellis Aune for discussions and help. We thank the BioFrontiers sequencing core at CU-Boulder. Libraries and shRNA clones were acquired from the functional genomics core at CU-Anschutz. Spinning disk confocal imaging was performed at the BioFrontiers Advanced Imaging Core supported by HHMI. FACS sorter and analyzer are supported by S10OD021601. This work was supported by the Searle Family Foundation (J.M.K) and the NIGMS (DP2GM123458, J.M.K).

## Competing interest statement

The authors have filed a provisional patent for cal-Seq.

## Supplementary Figures

**Supplementary Figure 1:**
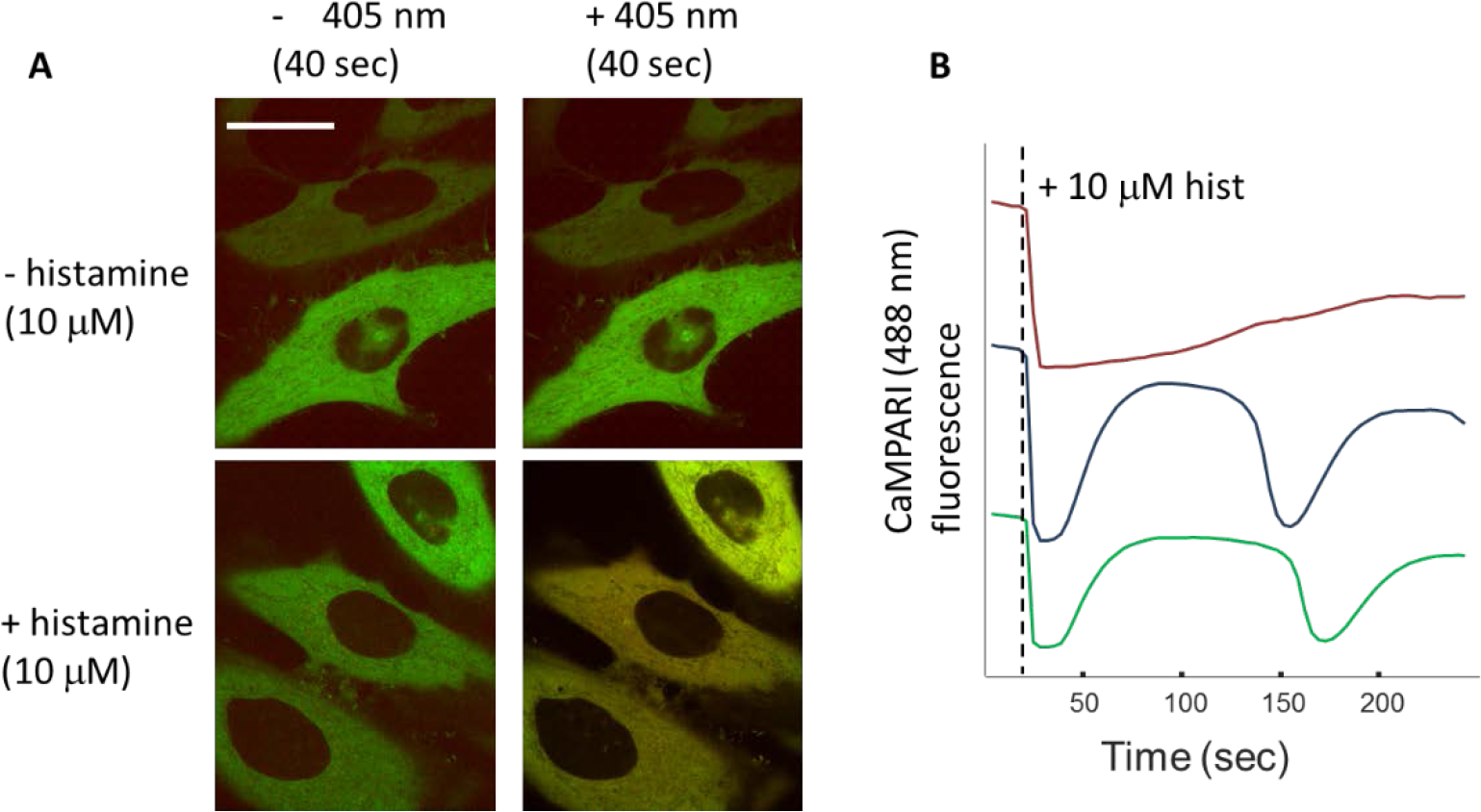
(A) HeLa cells expressing CaMPARI were fluorescent using a 488 nm excitation with the fluorescence confined to the cytoplasm (left column). Upon exposure to 40 seconds of 405 nm light, cells treated with histamine increased fluorescence excited by 561 nm light (bottom), whereas untreated cells remained green (top). Scale bar is 10 μm. (B) CaMPARI can act as a real time calcium indicator. HeLa cells expressing CaMPARI were imaged with 488 nm light and were stimulated with 100 μM histamine. The green fluorescence decreased indicating increased cytoplasmic calcium, similar to previous reports.

**Supplementary Figure 2:**
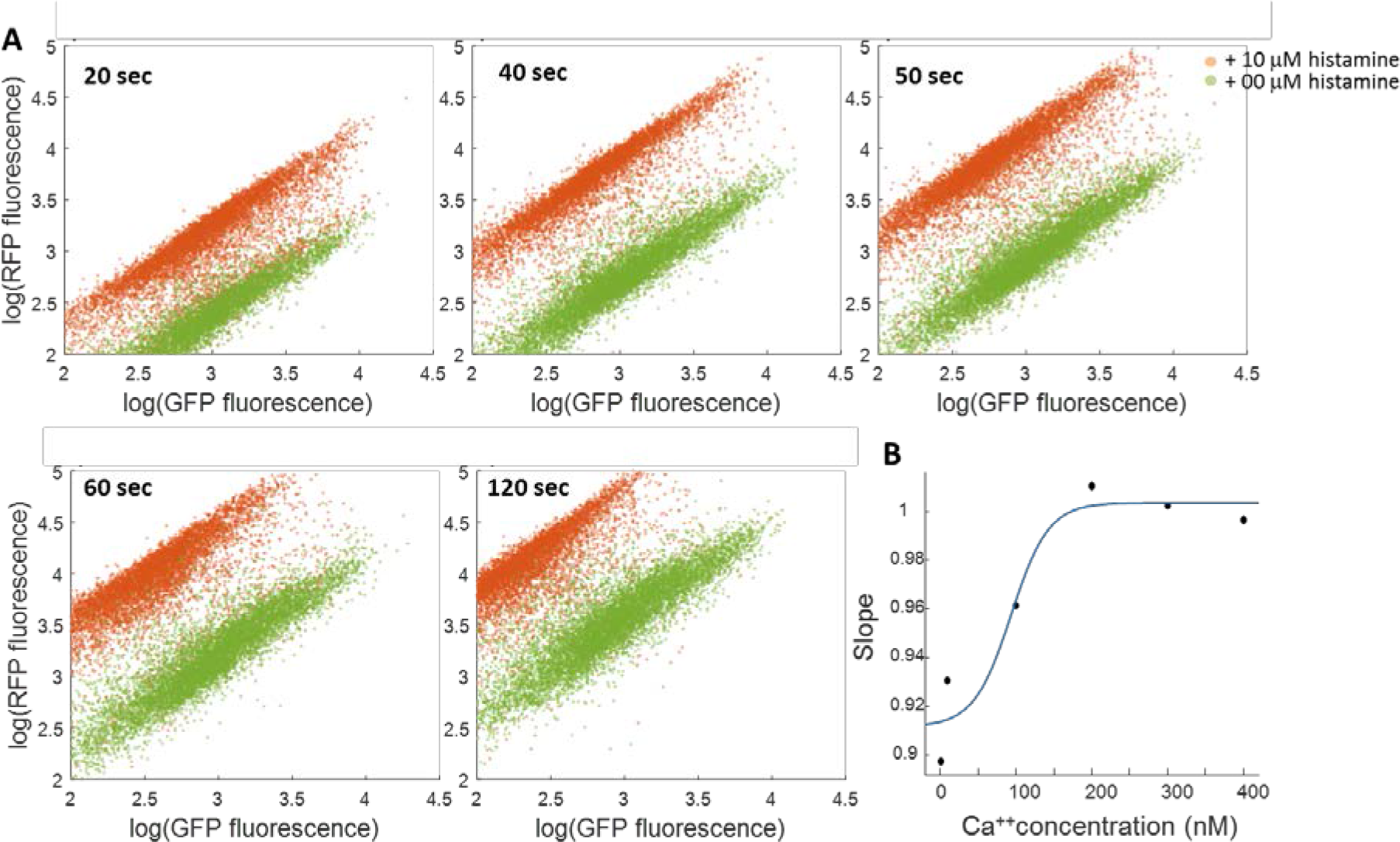
(A) Cytometry data showing the effect of increased 405 nm exposure time on CaMPARI expressing cells treated with 0 μM (green) or 10 μM (red) histamine. The largest separation between treated and untreated populations occurred at 40 seconds, and we used that timing for all additional experiments. (B) Kd calculations from the ionomycin cytometry data. The slopes of the best fit lines were fit to a hill curve which yielded a Kd = 124 nM calcium.

**Supplementary Figure 3:**
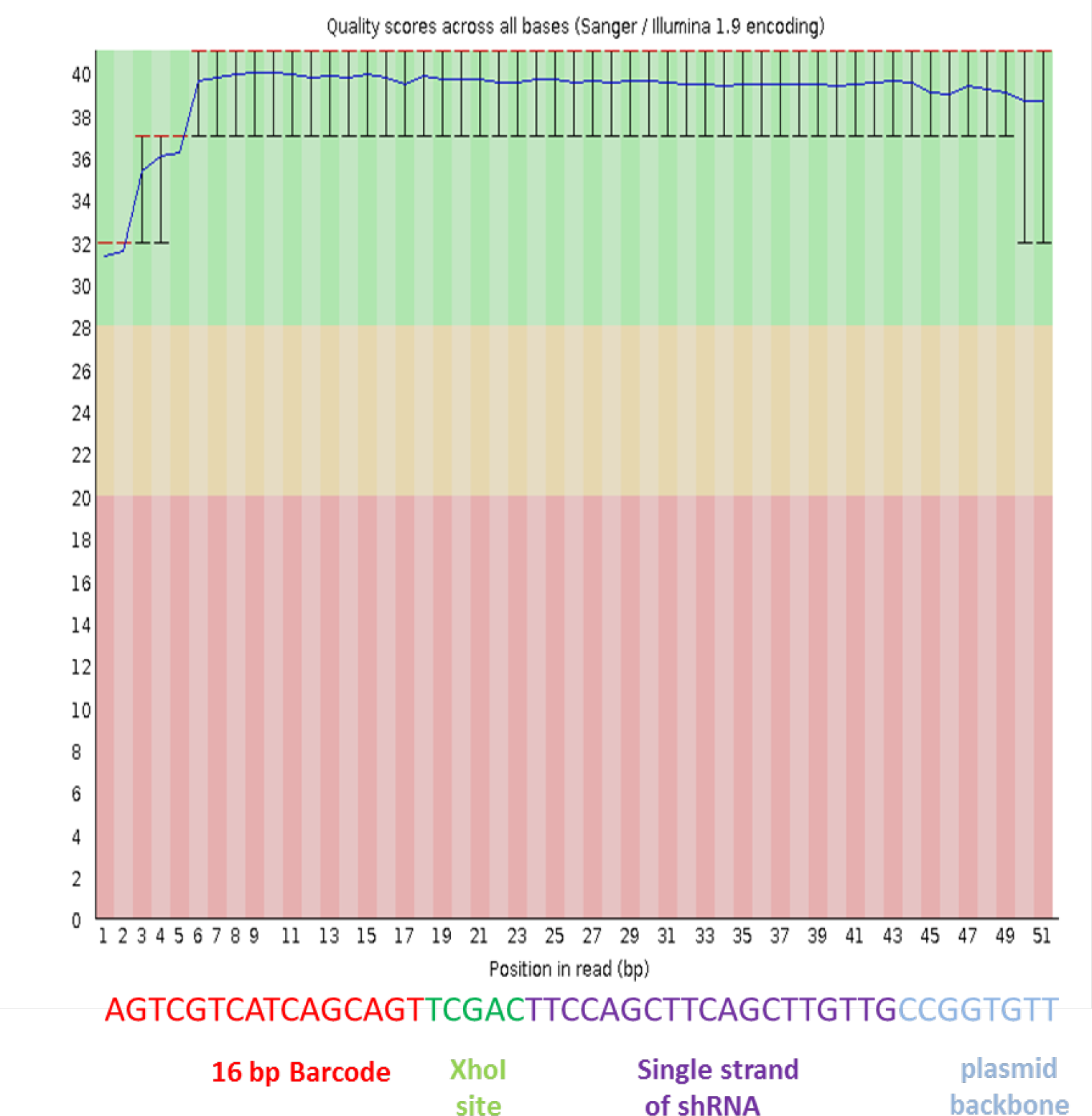
Read quality from the NextSeq was high across all positions of the shRNA tag. For each run, we spiked in 20% phiX to increase library diversity. The typical number of reads that matched to sequences in the TRC library was ~240M.

**Supplementary Figure 4:**
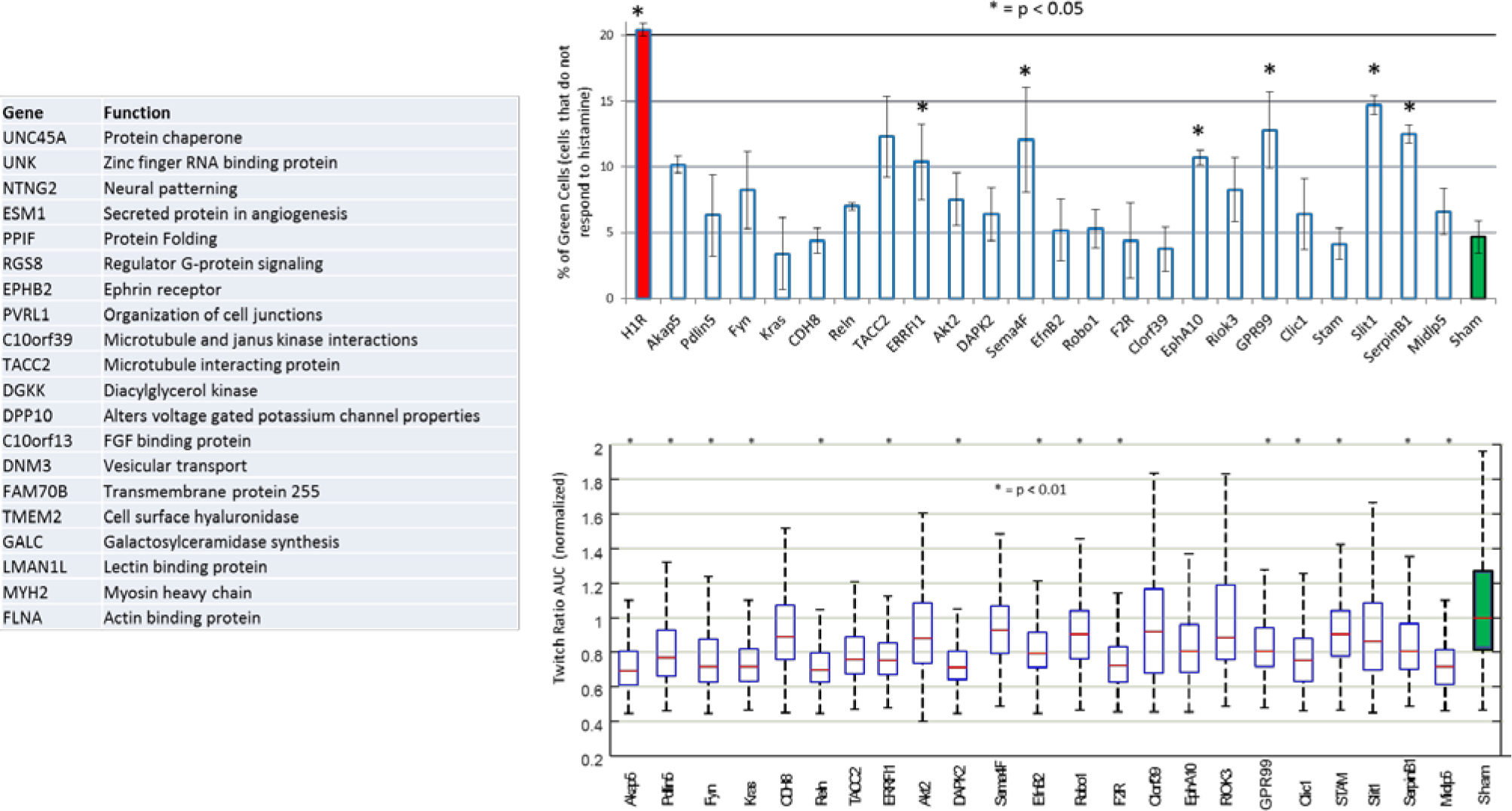
(A) List of the top 25 genes enriched in the green population as called by DeSeq analysis. (B) Cytometry data showing the fraction of green cells as compared to a sham when infected with single clone knockdowns identified in the enriched sequence. * represents a p-value < 0.05. (C) Calcium area-under-the-curve measurements for the top 25 hits using the real time indicator, Twitch2B. Cells were infected with lentivirus containing the single clone knockdown and imaged during histamine addition. Each bar chart represents 2 biological replicates with < 50 cells per field of view. * represents a p-value < 0.01.

**Supplementary Figure 5:**
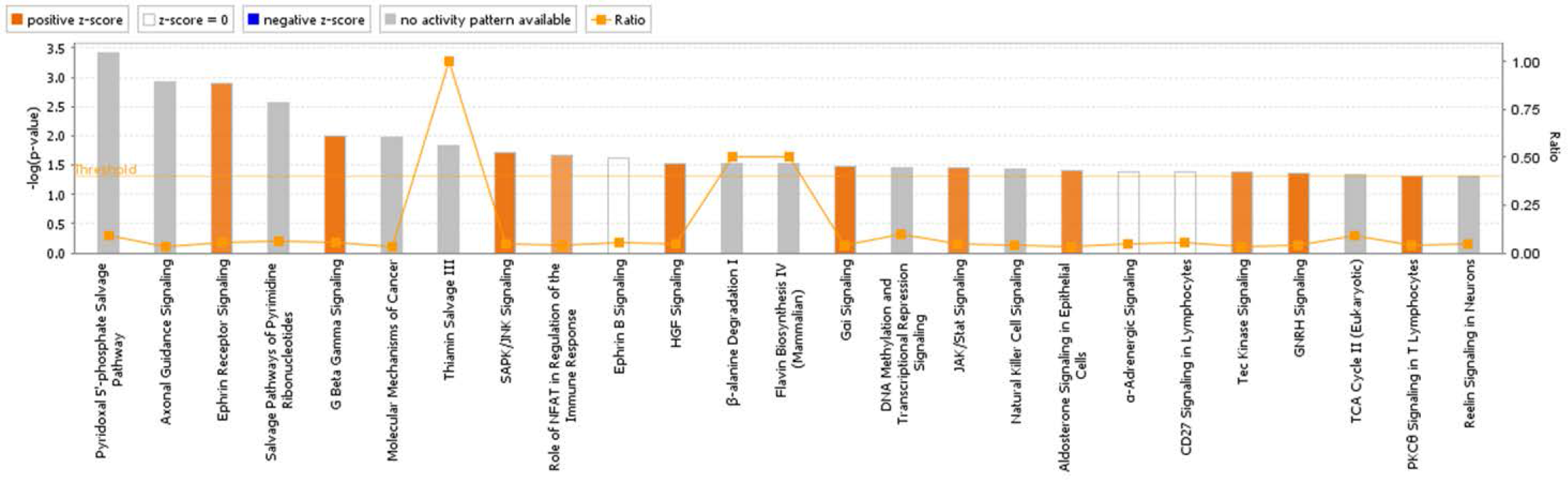
The top pathways identified from the DeSeq genes as identified by the Ingenuity Pathway Analysis software.

**Supplementary Figure 6:**
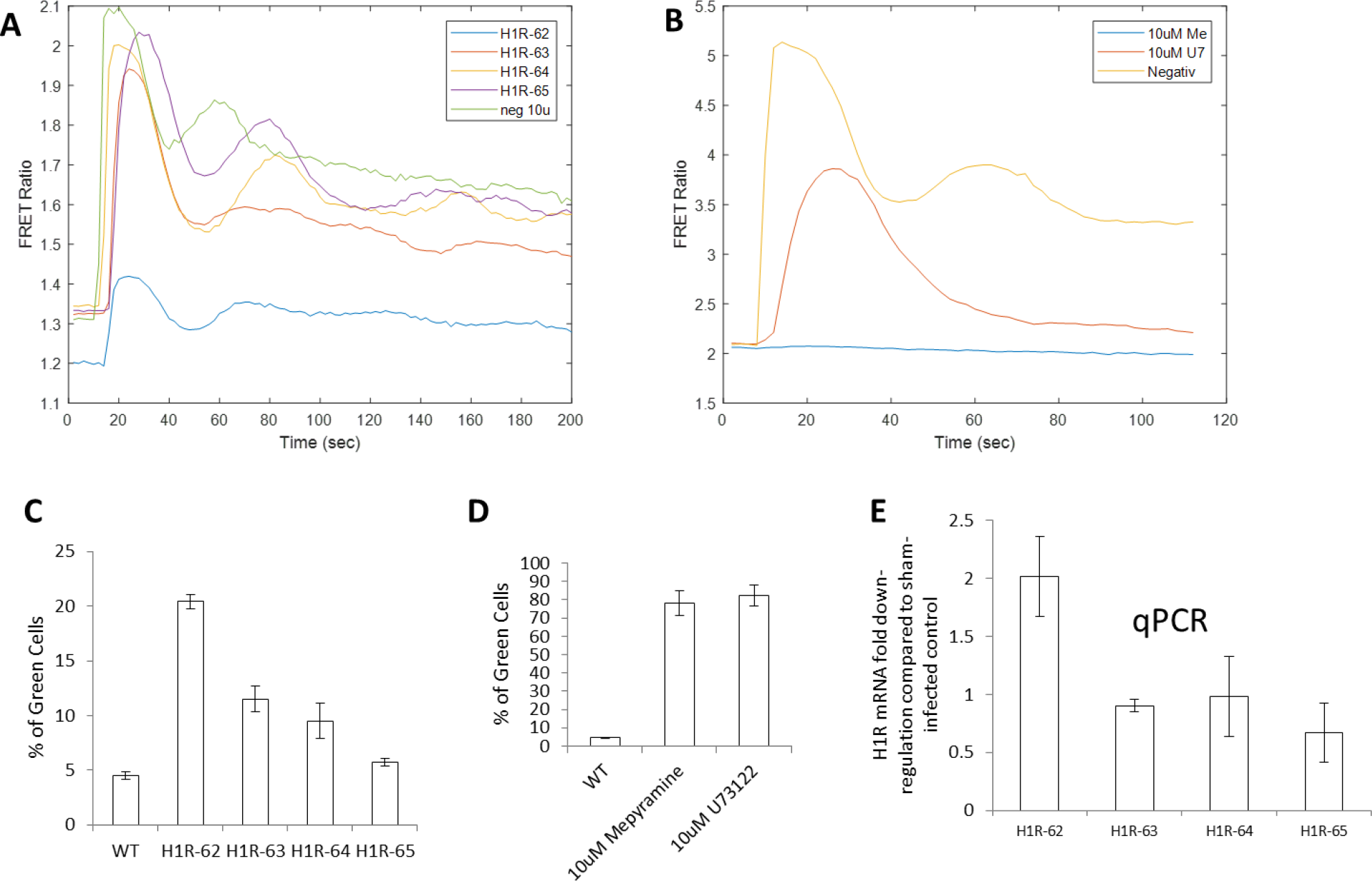
Chemical and genetic manipulation of the H1R receptor. (A) The TRC library has 4 shRNA targets against H1R, but only a single strand showed decreased calcium influx compared to a sham negative control as measured with Twitch. (B) Twitch measurements showing the effects of Mepyramine (H1R inhibitor) and U73122 (Phospholipase-C inhibitor). Mepyramine blocks all calcium influx upon histamine addition. (C) Cytometry measurements using CaMPARI of individual shRNA knockdowns. Only H1R-62 had a significantly increased fraction of green cells, however, it did not appear in our hit list. (D) Cytometry measurements of the H1R antagonists Mepyramine and U73122. (E) QPCR data revealed only a 2-fold knockdown of the H1R receptor using the H1R-62 shRNA which may explain why it did not appear in our screen.

**Supplementary Figure 7:**
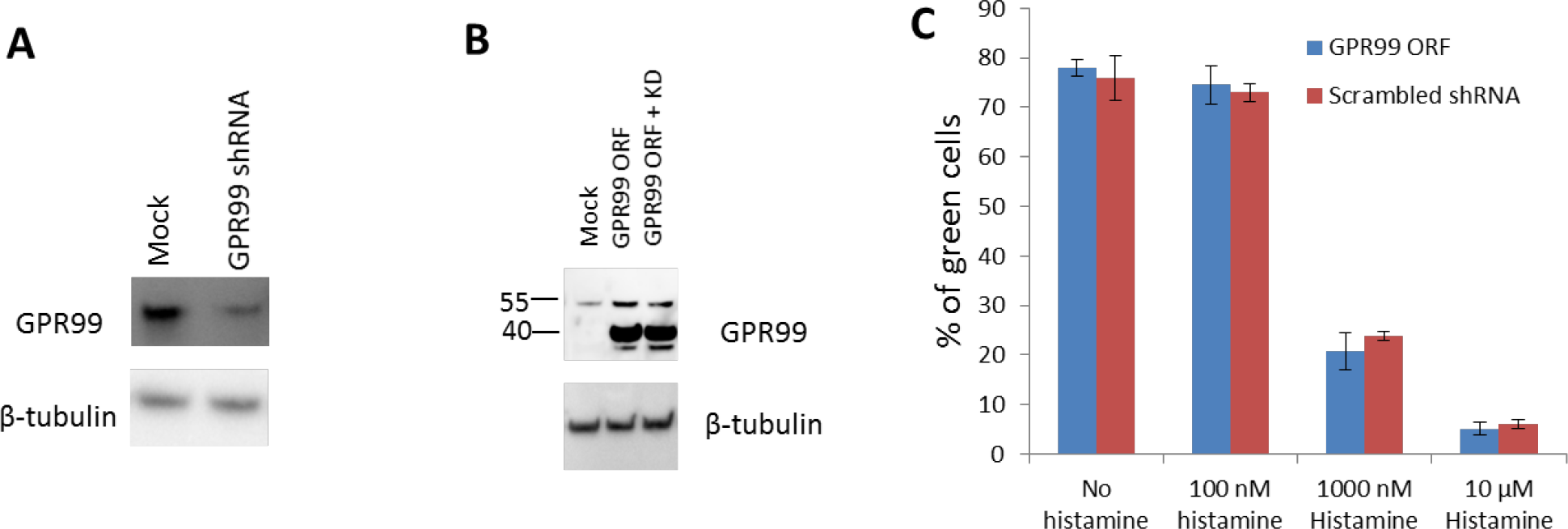
GPR99 ORF expression does not rescue the phenotype. (A) A western blot showing knockdown in HeLa cells with the shRNA. (B) Western blot showing the ORF expression. A second dominant band at 40 kD is visible only in the ORF expression, suggesting there are post-translational modifications on the endogenous receptor absent from the ORF construct. The shRNA did not reduce ORF expression. (C) Cytometry measurements showing histamine response of the GPR99 ORF. Overexpression of GPR99 did not alter the calcium influx from histamine stimulation as compared to a sham shRNA.

**Supplementary Figure 8:**
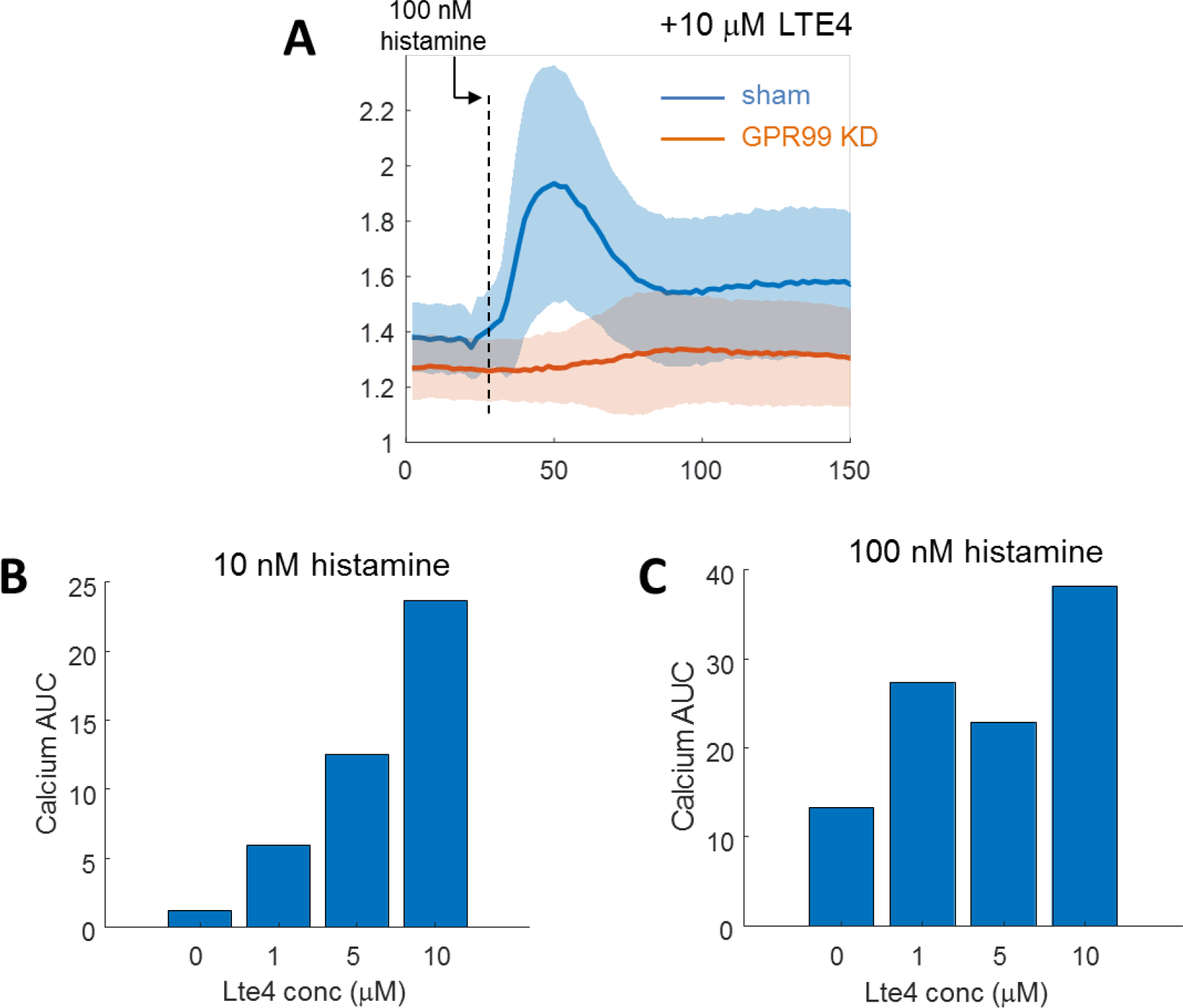
Increasing Lte4 concentration increases histamine induced calcium flux dependent on the presence of GPR99. (A) Pre-treatment of HeLa cells with Lte4 caused an increased calcium response in the sham shRNA (blue) but not in a GPR99 knockdown (red). The solid line shows population mean and the shaded area shows the standard deviation. (B and C). Pre-treatment of HeLa cells caused increased calcium responses with addition of 10 nM (B) or 100 nM (C) histamine. Calcium influx was measured with Twitch by taking the AUC of the first 40 seconds after histamine addition.

**Supplementary Figure 9:**
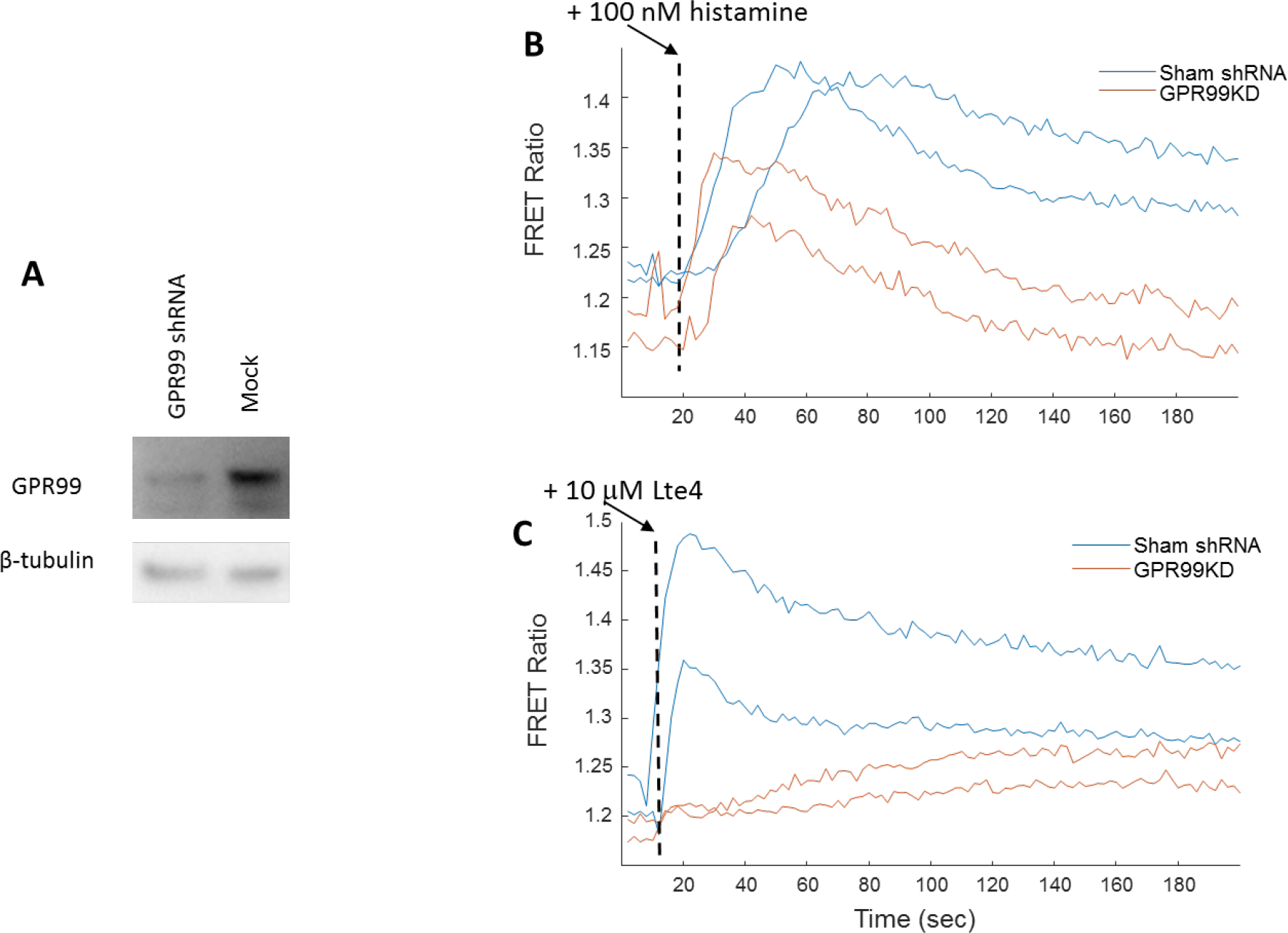
Beas2B epithelial cells also have histamine induced calcium influx dependent on GPR99 and Lte4. (A) Western blot showing a reduction of protein expression in Beas2B cells with the shRNA. (B) Histamine induced calcium flux as measured by Twitch is decreased in a GPR99 knockdown (red) as compared to a sham shRNA (blue). Each line represents the population mean of 1 biological replicate (> 60 cells). (C) Lte4 addition induced increased cytoplasmic calcium in sham (blue) but not GPR99 knockdown (red) cells.

**Supplementary Figure 10:**
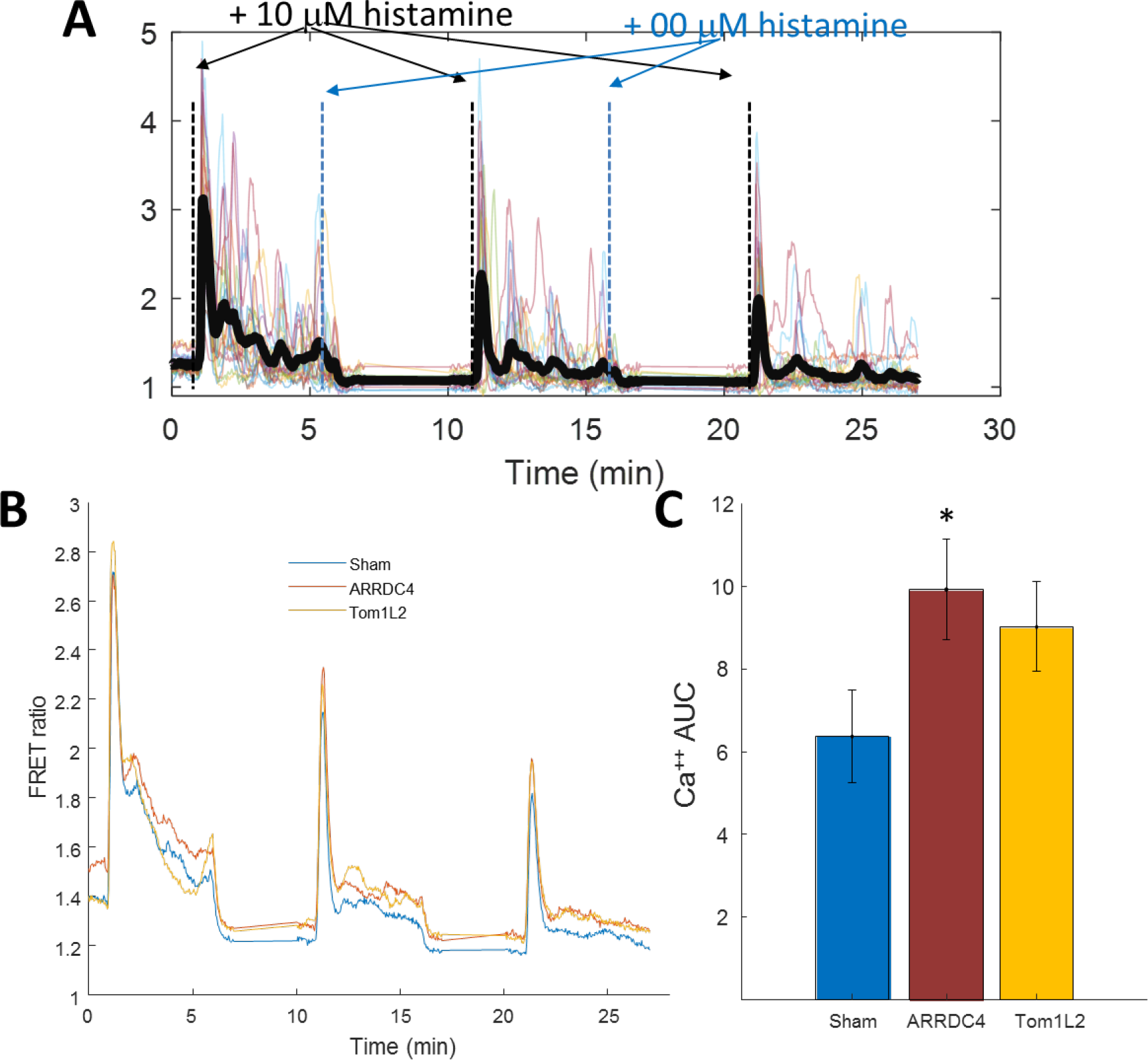
(A) Histamine induced desensitization protocol. HeLa cells were measured with Twitch upon repeated exposure and washout of 10 μM histamine (black and blue dashed lines, respectively). Imaging was paused after the washout to minimize photobleaching. The black line represents the population average. Semi-transparent colored lines represent individual cells to highlight the diversity in dynamics. (B) Average traces of 3 biological replicates showing the desensitization response of a sham (blue), ARRDC4 knockdown (red), and Tom1L2 knockdown (yellow). (C) Area under the curve measurements (40 seconds total time) after 2.5 minutes of histamine addition on the third stimulation for sham (blue), ARRDC4 knockdown (red), and Tom1L2 knockdown (yellow). * represents a p-value < 0.05 in a student t-test.

**Supplementary Figure 11:**
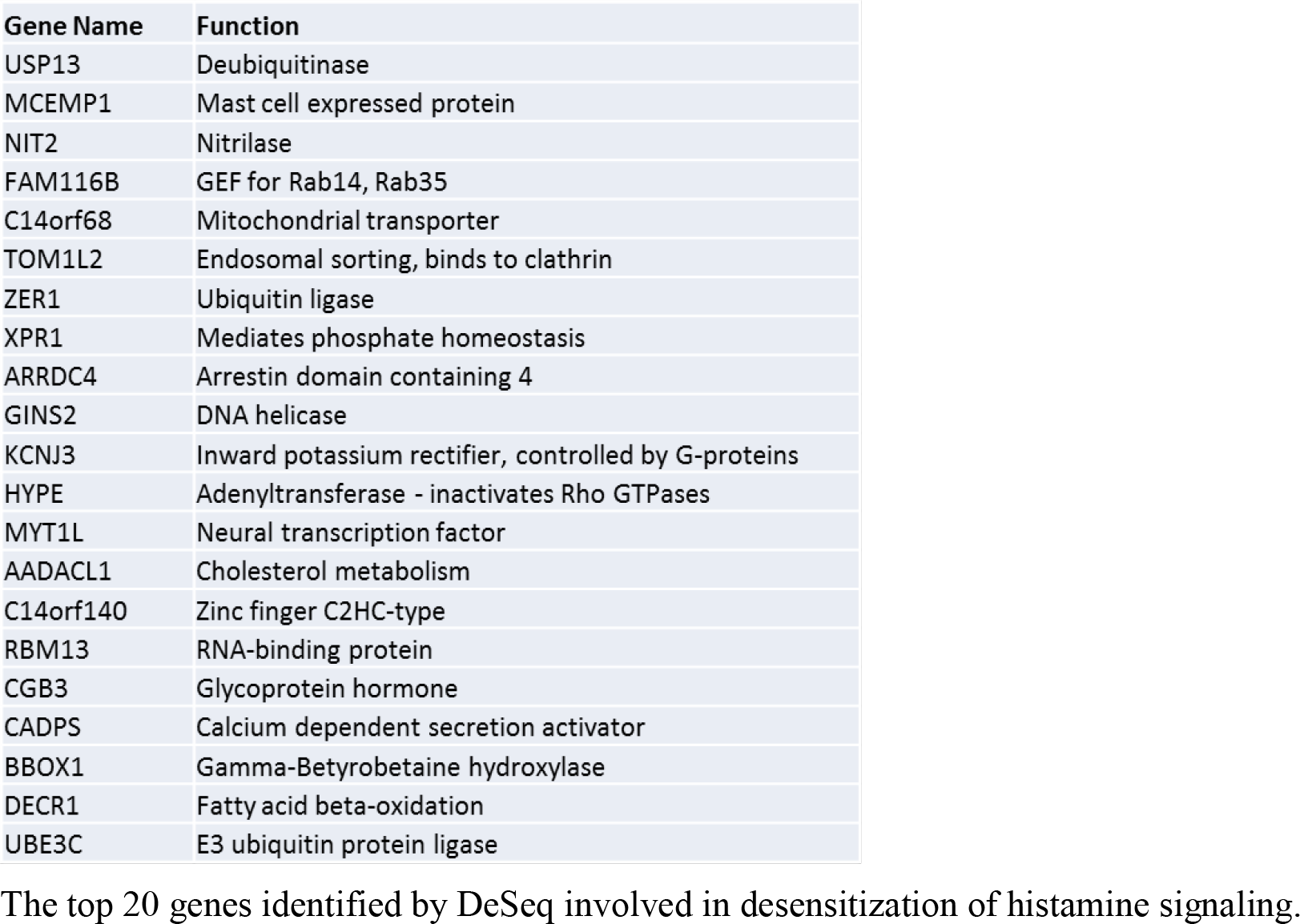
The top 20 genes identified by DeSeq involved in desensitization of histamine signaling.

## References (20)

1. Clapham, D. E., Calcium signaling. Cell 131, 1047–58 (2007).

2. Missiaen, L., et al. Abnormal intracellular Ca2+homeostasis and disease. Cell Calcium 28, 1–21 (2000).

3. Morgens, D. W., Deans, R. M., Li, A., & Bassik, M. C., Systematic comparison of CRISPR/Cas9 and RNAi screens for essential genes. Nat. Biotechnol. 34, 634–636 (2016).

4. Sharma, S., et al. An siRNA screen for NFAT activation identifies septins as coordinators of store-operated Ca2+ entry. Nature 499, 238–242 (2013).

5. Schmunk, G., et al. High-throughput screen detects calcium signaling dysfunction in typical sporadic autism spectrum disorder. Sci. Rep. 7, 40740 (2017).

6. Fosque, B. F., et al. Labeling of active neural circuits in vivo with designed calcium integrators. Science (80-.). 347, 755–760 (2015).

7. Root, D. E., Hacohen, N., Hahn, W. C., Lander, E. S., & Sabatini, D. M., Genome-scale loss-of-function screening with a lentiviral RNAi library. Nat. Methods 3, 715–719 (2006).

8. Anders, S., & Huber, W., Differential expression analysis for sequence count data. Genome Biol. 11, R106 (2010).

9. Hart, T., Brown, K. R., Sircoulomb, F., Rottapel, R., & Moffat, J., Measuring error rates in genomic perturbation screens: gold standards for human functional genomics. Mol. Syst. Biol. 10, 733 (2014).

10. Thestrup, T., et al. Optimized ratiometric calcium sensors for functional in vivo imaging of neurons and T lymphocytes. Nat. Methods 11, 175–82 (2014).

11. Foskett, J. K., White, C., Cheung, K.-H., & Mak, D.-O. D., Inositol trisphosphate receptor Ca2+ release channels. Physiol. Rev. 87, 593–658 (2007).

12. Hogan, P. G., Chen, L., Nardone, J., & Rao, A., Transcriptional regulation by calcium, calcineurin, and NFAT. Genes Dev. 17, 2205–32 (2003).

13. Sutherland, D. J., Pujic, Z., & Goodhill, G. J., Calcium signaling in axon guidance. Trends Neurosci. 37, 424–432 (2014).

14. Mirakaj, V., & Rosenberger, P., Immunomodulatory Functions of Neuronal Guidance Proteins. Trends Immunol. 38, 444–456 (2017).

15. Kanaoka, Y., Maekawa, A., & Austen, K. F., Identification of GPR99 Protein as a Potential Third Cysteinyl Leukotriene Receptor with a Preference for Leukotriene E 4 Ligand. J. Biol. Chem. 288, 10967–10972 (2013).

16. Bankova, L. G., et al. Leukotriene E 4 elicits respiratory epithelial cell mucin release through the G-protein-coupled receptor, GPR99. Proc. Natl. Acad. Sci. 113, 6242–6247 (2016).

17. Wittenberger, T., et al. GPR99, a new G protein-coupled receptor with homology to a new subgroup of nucleotide receptors. BMC Genomics 3, 17 (2002).

18. Bloemers, S. M., et al. Sensitization of the histamine H1 receptor by increased ligand affinity. J. Biol. Chem. 273, 2249–55 (1998).

19. Xatzipsalti, M., & Papadopoulos, N. G., Cellular and animals models for rhinovirus infection in asthma. Contrib. Microbiol. 14, 33–41 (2007).

20. Bartho, L., & Benko, R., Should antihistamines be re-considered as antiasthmatic drugs as adjuvants to anti-leukotrienes? Eur. J. Pharmacol. 701, 181–4 (2013).

21. Austen, K. F., Maekawa, A., Kanaoka, Y., & Boyce, J. A., The leukotriene E4 puzzle: Finding the missing pieces and revealing the pathobiologic implications. J. Allergy Clin. Immunol. 124, 406–414 (2009).

22. Freedman, N. J., & Lefkowitz, R. J., Desensitization of G protein-coupled receptors. Recent Prog. Horm. Res. 51, 319–51-3 (1996).

23. Smit, M. J., et al. Short-term desensitization of the histamine H1 receptor in human HeLa cells: involvement of protein kinase C dependent and independent pathways. Br. J. Pharmacol. 107, 448–55 (1992).

24. Sullivan, K. D., et al.Trisomy 21 consistently activates the interferon response. Elife 5, e16220 (2016).

